# Hyperbolic graph embedding of MEG brain networks to study brain alterations in individuals with subjective cognitive decline

**DOI:** 10.1101/2023.10.23.563643

**Authors:** Cole Baker, Isabel Suárez-Méndez, Grace Smith, Elisabeth B. Marsh, Michael Funke, John C. Mosher, Fernando Maestú, Mengjia Xu, Dimitrios Pantazis

**Affiliations:** Mc-Govern Institute for Brain Research, Massachusetts Institute of Technology, Cambridge, MA 02139, USA; Department of Data Science, Ying Wu College of Computing, New Jersey Institute of Technology, Newark, NJ 07102, USA; Department of Experimental Psychology, Complutense University of Madrid, Madrid 28040, Spain; Department of Neurology, Johns Hopkins University School of Medicine, Baltimore, MD 21205, USA; Department of Neurology, McGovern Medical School, UTHealth Houston, Houston, TX 77030, USA

**Author notes:** (*Corresponding authors:* Dimitrios Pantazis and Mengjia Xu) and).

**Keywords:** Alzheimer’s disease, subjective cognitive decline, magnetoencephalography, brain networks, hyperbolic space, graph embedding

## Abstract

An expansive area of research focuses on discerning patterns of alterations in functional brain networks from the early stages of Alzheimer’s disease, even at the subjective cognitive decline (SCD) stage. Here, we developed a novel hyperbolic MEG brain network embedding framework for transforming high-dimensional complex MEG brain networks into lower-dimensional hyperbolic representations. Using this model, we computed hyperbolic embeddings of the MEG brain networks of two distinct participant groups: individuals with SCD and healthy controls. We demonstrated that these embeddings preserve both local and global geometric information, presenting reduced distortion compared to rival models, even when brain networks are mapped into low-dimensional spaces. In addition, our findings showed that the hyperbolic embeddings encompass unique SCD-related information that improves the discriminatory power above and beyond that of connectivity features alone. Notably, we introduced a unique metric—the radius of the node embeddings—which effectively proxies the hierarchical organization of the brain. Using this metric, we identified subtle hierarchy organizational differences between the two participant groups, suggesting increased hierarchy in the dorsal attention, frontoparietal, and ventral attention subnetworks among the SCD group. Last, we assessed the correlation between these hierarchical variations and cognitive assessment scores, revealing associations with diminished performance across multiple cognitive evaluations in the SCD group. Overall, this study presents the first evaluation of hyperbolic embeddings of MEG brain networks, offering novel insights into brain organization, cognitive decline, and potential diagnostic avenues of Alzheimer’s disease.

## I. Introduction

Alzheimer’s Disease (AD) has emerged as a global public health imperative, particularly among older populations. AD, characterized by a gradual and relentless neurodegenerative process, entails a protracted *asymptomatic* preclinical phase preceding the onset of mild cognitive impairment (MCI) and profound cognitive deterioration [1]. Currently, the impact of AD is palpable, affecting 6.5 million individuals in the United States alone, with projections indicating a staggering surge to 14 million by 2060 [2].

A rising phenomenon, subjective cognitive decline (SCD), is garnering attention as an emerging self-reported condition. It encapsulates the personal perception of cognitive decline, not necessarily paralleled by objective diminishment on standardized evaluations [3]. Increasing evidence shows that SCD may be a precursor to the early stages of Alzheimer’s disease (e.g., MCI) and related dementias [4]–[6]. Remarkably, individuals with SCD are five times more likely to progress to early stage of AD (e.g., MCI) than people without SCD [7]. Hence, detection and understanding of the key neural signatures and biomarkers of SCD at the presymptomatic stage of AD play an important role in developing preventative cognitive interventions, identifying pharmaceutical targets, and monitoring pathological progression in the preclinical stages of AD.

The two major histopathological hallmarks of AD are extracellular amyloid-*β* plaques and intracellular neurofibrillary tangles, stemming from the phosphorylation of the Tau protein [8]. Amyloid deposition has the detrimental effect of impairing normal inter-neuronal connectivity [9], while the presence of Tau proteins leads to the disruption of axonal microtubule organization [10]. This sequence of events culminates in the progressive deterioration and pruning of synaptic connections, disrupting communication both within and between brain regions [11], and giving rise to distinctive patterns of alterations in functional connectivity within large-scale brain systems. Ongoing research efforts aim to discern such patterns from the early stages of AD, even at the SCD stage, using non-invasive neuroimaging methods, including functional magnetic resonance imaging (fMRI) [12], [13], positron emission tomography (PET) [14], and magnetoencephalography (MEG) [15].

Among these methods, MEG offers excellent temporal resolution that allows characterization of the subtle brain changes associated with AD, a major advantage over current biomarkers [11], [16]. This is especially important because, unlike methods that rely on metabolic responses (fMRI, FDG-PET), functional brain networks can also be characterized in the frequency domain [17], [18]. Moreover, MEG captures the fields produced by intraneuronal currents, providing a more direct index of neuronal activity than biomarkers that measure haemodynamic responses. Thus, MEG holds great promise in detecting synaptic dysfunctions in the brains of those exhibiting varying degrees of cognitive decline, especially in characterizing the subtle neural activity changes between SCD and healthy controls. However, SCD studies with MEG are still scarce, with most existing MEG-based SCD studies heavily relying on statistical analyses with traditional ad-hoc features within pre-selected brain subnetworks, such as the default mode network (DMN) and dorsal attention network (DAN), for discriminating between SCD, mild cognitive impairment (MCI), and AD groups [19], [20].

Recently, neural network models have shown great potential in *embedding* complex human brain networks into latent spaces where nodes are represented as low-dimensional vectors [21] or probabilistic density functions [13], [15], [22]. The original graph topological properties and node similarity are maximally preserved in these latent spaces. Moreover, the derived node embeddings can be readily and efficiently applied to diverse downstream tasks, including link prediction, node classification, and community detection [23].

However, the quality of graph embeddings critically depends on whether the geometry of the embedding space matches the underlying structure of the graph [24]. The prevalent approach in graph representation learning involves graph convolutional neural networks that embed the nodes of the graph into points in the *Euclidean space*. But the Euclidean geometry has limited representational capacity and high distortion when embedding brain networks. This is because brain networks are scale-free graphs with a tree-like hierarchical structure, where the number of nodes grows *exponentially* as we move away from higher towards lower hierarchy regions. This substantially exceeds the *polynomial* expanding space capacity of the Euclidean space, resulting in embeddings with high distortion [25], [26]. To mitigate this problem, a viable solution is to increase the dimension of the Euclidean embedding space. However, this remedy leads to complex models with reduced generalization capacity, higher computational costs, and increased memory requirements.

In contrast, the *hyperbolic space*, characterized by negative curvature, offers a direct solution to the exponential expansion of the number of nodes in scale-free brain graphs [27]–[29]. This is because the hyperbolic geometry is characterized by an exponential growth of space as we move away from the center, mirroring the exponential growth of brain networks. The merits of this approach for embedding brain networks are as follows [30]:

- It yields embeddings with minimal distortion, thereby preserving both local and global geometric information;
- Due to the minimal distortion, brain networks can be mapped into low-dimensional spaces, facilitating downstream tasks (such as SCD classification here)
- The neural network model has improved generalization capacity, less complexity, and lower training data requirements.
- Importantly, the resulting embeddings possess an intuitive interpretation property. Higher hierarchy brain nodes map closer to the geometry’s center, while lower hierarchy nodes map towards the periphery. This representational characteristic opens new avenues for novel insights regarding the hierarchical organization intrinsic to brain networks, a key focal point of this study.

Here, we developed a novel hyperbolic MEG brain network embedding framework for transforming high-dimensional complex MEG brain networks into lower-dimensional hyperbolic representations. Our approach involved the design and validation of a new hyperbolic model build upon the architecture of the hyperbolic graph convolutional network (HGCN) model [25]. Using this model, we computed hyperbolic embeddings of the MEG brain networks of two distinct participant groups: individuals with SCD and healthy controls. To assess the quality and utility of our embeddings, we compared the network distortion against competing models in both Euclidean and manifold-based hyperbolic embeddings derived from a shallow Poincaré embedding model [27]. We then leveraged these embeddings to differentiate between SCD and healthy participants, thereby evaluating their discriminatory power. Notably, we introduced a unique metric—the radius of the node embeddings—which effectively proxies the hierarchical organization of the brain. We used this metric to identify brain subnetworks characterized by subtle hierarchy organizational differences between the two participant groups. Finally, our investigation assessed the correlation between these hierarchical variations and cognitive assessment scores, thereby providing insights into the potential cognitive implications of the identified brain hierarchical differences.

## II Methods

### A Participants and cognitive intervention training

The cohort for this study comprised 146 participants recruited from the Centre for Prevention of Cognitive Impairment (Madrid Salud), the Faculty of Psychology of the Complutense University of Madrid (UCM), and the hospital Clinico San Carlos (HCSC) in Madrid, Spain between January 2014 and December 2015 [19], [31]. All participants were between 65-80 years old, right-handed, and Spanish natives. This study received ethical approval from the Ethics Committee of the HCSC, and prior to their inclusion, every participant provided informed consent.

During the initial screening phase, all participants underwent a battery of cognitive assessments, as elaborated in [31]. They were subsequently categorized into either the Subjective Cognitive Decline (SCD) or healthy control (HC) groups, adhering to the research criteria outlined by the Subjective Cognitive Decline Initiative (SCD-I) [3]. Specifically, the diagnosis of SCD was established with the exclusion of potential confounding factors associated with cognitive impairment, including psychiatric or neurological diseases and mental disorders, drug use, and evidence of infection. The assessment followed a set of specific criteria, as outlined below: (1) Participants reported persistent cognitive concerns, primarily related to memory, during interviews conducted by experienced clinicians. (2) They demonstrated cognitive performance within the normal range on standardized tests capable of distinguishing individuals with MCI and prodromal AD. (3) Participants expressed that their cognitive decline had a discernible impact on their daily activities. (4) They actively sought medical consultation regarding their cognitive issues. (5) Individuals diagnosed with SCD were aged 60 years or older at the onset of these cognitive concerns, and these concerns had arisen within the past 5 years. Additionally, the cognitive decline was corroborated by reliable informants.

The cognitive assessments, along with MEG and MRI recordings, were conducted both at the time of enrollment and again 6 months later, as part of a 6-month training intervention study designed at the Memory Training Unit of the City Council of Madrid, as detailed in [31]. Participants were randomly divided into trained and non-trained groups. The trained group underwent a 30-session intervention (28 regular sessions and two maintenance booster sessions), lasting 90 minutes each. These sessions occurred three times a week in the morning and were conducted by professionals. They encompassed various factors, including cognitive stimulation, learning of cognitive strategies, interventions in daily living performance, and analysis of metamemory. Additionally, they promoted a healthy lifestyle through activities like proper nutrition, physical exercise, social engagement, and leisure activities.

The eligible participants were further narrowed by removing 34 patients who did not attend the second MEG session and 22 patients whose scans presented technical reconstruction issues. Among the remaining 90 patients, the trained group comprised 46 subjects (22 HC and 24 SCD), and the non-trained group comprised 44 subjects (19 HC and 25 SCD subjects). Demographics for the 90 participants are described in the Table I.

**TABLE I.**
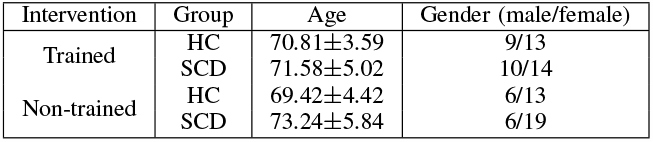
Demographic characteristics of all participants.

| Intervention | Group | Age | Gender (male/female) |
| --- | --- | --- | --- |
| Trained | HC | 70.81±3.59 | 9/13 |
|  | SCD | 71.58±5.02 | 10/14 |
| Non-trained | HC | 69.42±4.42 | 6/13 |
|  | SCD | 73.24±5.84 | 6/19 |

### B Data acquisition and pre-processing

Four minutes of resting-state MEG data was collected for each participant using a 306-channel Vectorview MEG system (Elekta AB, Stockholm, Sweden) at the Laboratory of Cognitive and Computational Neuroscience within the Centre for Biomedical Technology (Madrid, Spain). The MEG recordings were conducted at a sampling rate of 1000 Hz, and an online anti-alias band-pass filter was applied, ranging from 0.1 to 330 Hz. To suppress external magnetic interference, recordings were then processed using the temporally extended signal space separation method [32]. Finally, we applied independent component analysis to remove EOG and EKG components from the data.

### C Source reconstruction and connectivity analysis

The MEG time courses were segmented into non-overlapping 4-second epochs containing artifact-free activity. The number of clean epochs did not differ across groups and conditions. The epochs were subsequently band-pass filtered in the alpha band (8-12 Hz). For source reconstruction, the source space comprised 1220 sources placed in a homogeneous grid of 1 cm using the Montreal Neurological Institute (MNI) template, which was converted to subject space by affine transformation. MEG sources were anatomically mapped to 90 regions of interest (ROI) based on the Automated Anatomical Labeling (AAL) atlas [33]. Forward modeling used a single-shell head model [34], defined by the inner skull boundary generated from individual T1-weighted MRI scans using the Fieldtrip toolbox. Source reconstruction was performed for each subject using a Linearly Constrained Minimum Variance (LCMV) beamformer [35]. Beamformer Filters were obtained using the computed lead field, the epoch-averaged covariance matrix, and a 1% regularization factor.

Functional connectivity between individual sources was estimated using the phase locking value (PLV), a metric that assesses source-to-source connectivity based on phase synchronization principles [34]. Subsequently, region-of-interest (ROI) connectivity was computed using Eq. 1 by averaging the pairwise PLV values across all source pairs belonging to the two ROIs:

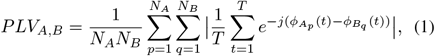

where 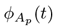 and 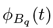 are the instantaneous phases of the signals *A*_*p*_(*t*) and *B*_*q*_(*t*) at time point *t, T* is the number of time points per epoch, *j* is the imaginary unit, *N*_*A*_ is the number of sources in region A, and *A*_*p*_ is the *p*-th source within that specific region.

This computation yielded two 90×90 connectivity matrices for each participant, during both the pre- and post-intervention recording sessions. Finally, to transform the connectivity matrices into binary graphs suitable for HGCN processing, we applied a threshold of 0.329 to the PLV values, retaining approximately 20% of the edges (Fig. 1).

**Fig. 1.**
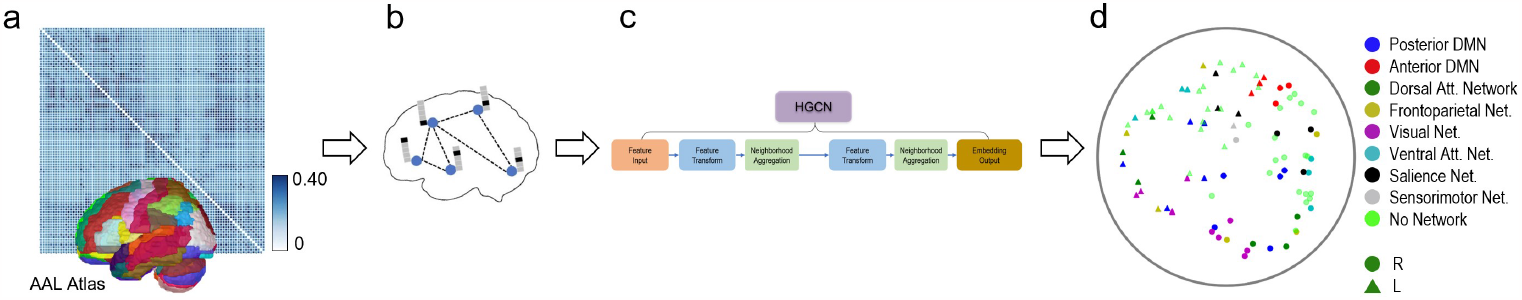
Workflow of the hyperbolic embedding framework for MEG brain networks. (a) MEG functional connectivity matrix calculated based on phase locking values across all 90 regions of the AAL brain atlas. (b) Illustrative visualization of the binary MEG brain network on the cortex, after applying a threshold of 0.329 to the phase locking values. Node attributes incorporate one-hot identity vectors. (c) Hyperbolic graph convolutional network. (d) Example embeddings of brain regions in the 2-dimensional space. Subnetwork and hemisphere are indicated by color and shape, respectively.

### D Assignment of brain regions to subnetworks

Brain subnetworks are composed of various brain regions that exhibit synchronized or coordinated activity when the brain is engaged in specific tasks or cognitive processes [36]. Studying these subnetworks can provide insights into brain function, cognitive processes, and neurological disorders [37].

Atlases delineating the ROIs within these subnetworks often deviate significantly from structural atlases, such as the AAL, because they are based on patterns of communication instead of physical structures in the brain. To assign brain regions to distinct subnetworks, we conducted a comprehensive evaluation of various subnetwork studies [38]–[41]. We then devised a procedure that considered the spatial proximity between each AAL anatomical region and the central coordinates of different functional ROIs. Using this procedure, we categorised the 90 ROIs into 8 distinct brain subnetworks: the posterior default mode network (pDMN), anterior default mode network (aDMN), dorsal attention network (DAN), frontoparietal network (FPN), visual network (VN), ventral attention network (VAN), salience network (SN), and sensorimotor network (SMN). Details regarding this assignment of ROIs to subnetworks are provided in the Supplementary Material.

### E Hyperbolic embedding of MEG brain networks

To study brain connectivity differences between the SCD and HC groups, we developed a hyperbolic embedding model based on the HGCN architecture. Using a hyperbolic space has the advantages of optimally preserving the topological properties of brain networks, and providing novel hierarchical information inherent in the embedding space.

#### 1) Hyperbolic geometry

Hyperbolic geometry is a non-Euclidean geometry that studies spaces of negative curvature, *K* < 0. In network science, hyperbolic geometry has gained attention for its ability to model hierarchical data. One of the most prevalent hyperbolic models is the Poincaré ball, where the hyperbolic space is represented as an *n*-dimensional open ball of fixed radius 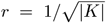. Distances between two points “cost” exponentially more as they reach the edge of the disk, such that distance to reach the edge itself is infinite. Shortest paths in this space are not straight lines, but arcs that bend closer to the origin to take advantage of the lower cost (Fig. 2a). The midpoint of that arc is analogous to a common parent node in the tree. Just as the number of child nodes in a branching tree grows exponentially with distance from the root, the continuous space for embeddings grows exponentially with distance from the origin. Networks with tree-like structures can be embedded in this space with fewer dimensions and with less distortion than in the Euclidean space (Fig. 2b) [27]–[29].

**Fig. 2.**
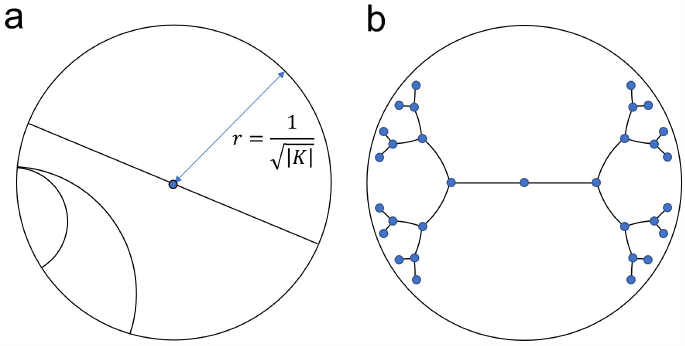
Poincaré ball embedding space. (a) Geodesic lines are arcs bent towards the origin. (b) Example Poincaré embedding of a binary tree.

Applying standard deep learning algorithms in hyperbolic space presents many challenges, as many standard operations are not obviously defined. To provide a foundational understanding, we offer a concise overview of essential equations in Poincaré ball and recommend relevant literature for a more comprehensive exploration [42]–[44].

One of the fundamental concepts in hyperbolic space is the geodesic distance. Minimizing geodesic distances between connected nodes is a common optimization objective:

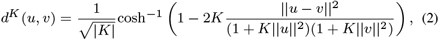

where the hyperbolic distance between points *u* and *v* depends on the curvature *K*.

Many operations necessary for hyperbolic neural networks are poorly defined or intractable in hyperbolic space. Following work in [42], most operations were formalized in a hybrid way, by transforming features between the hyperbolic space and a tangent space, which is a Euclidean subspace at the origin of the hyperbolic model, and performing the operations within the tangent space. Transportation between spaces is achieved via logarithmic and exponential maps.

The logarithmic map, denoted as *log*^*K*^, transforms a point *x* from a hyperbolic space with curvature *K* into its Euclidean projection. The desired function *f* is then applied within the Euclidean space, and the output of this function is transported back to the hyperbolic space using the exponential map, designated as *exp*^*K*^. All together:

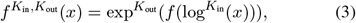

where *K*_in_ and *K*_out_ are the curvatures of the input and output hyperbolic spaces, respectively. The logarithmic and exponential maps allow for arbitrary functionality in the hyperbolic space, but introducing numerical errors and should be avoided when mathematically feasible.

Using this approach, the work in [42] derived close-form solutions in the hyperbolic space for matrix-vector multiplication and bias addition. For brevity, we present the symbolic notations:

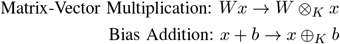

#### 2) Hyperbolic Graph Convolutional Network

The HGCN is a generalization of GCN to hyperbolic space [25]. The model takes as input a graph, in the form of a binary adjacency matrix, and node input features, and outputs hyperbolic embeddings suitable for downstream tasks, such as link prediction or graph classification. Our HGCN model architecture is illustrated in Fig. 3.

**Fig. 3.**
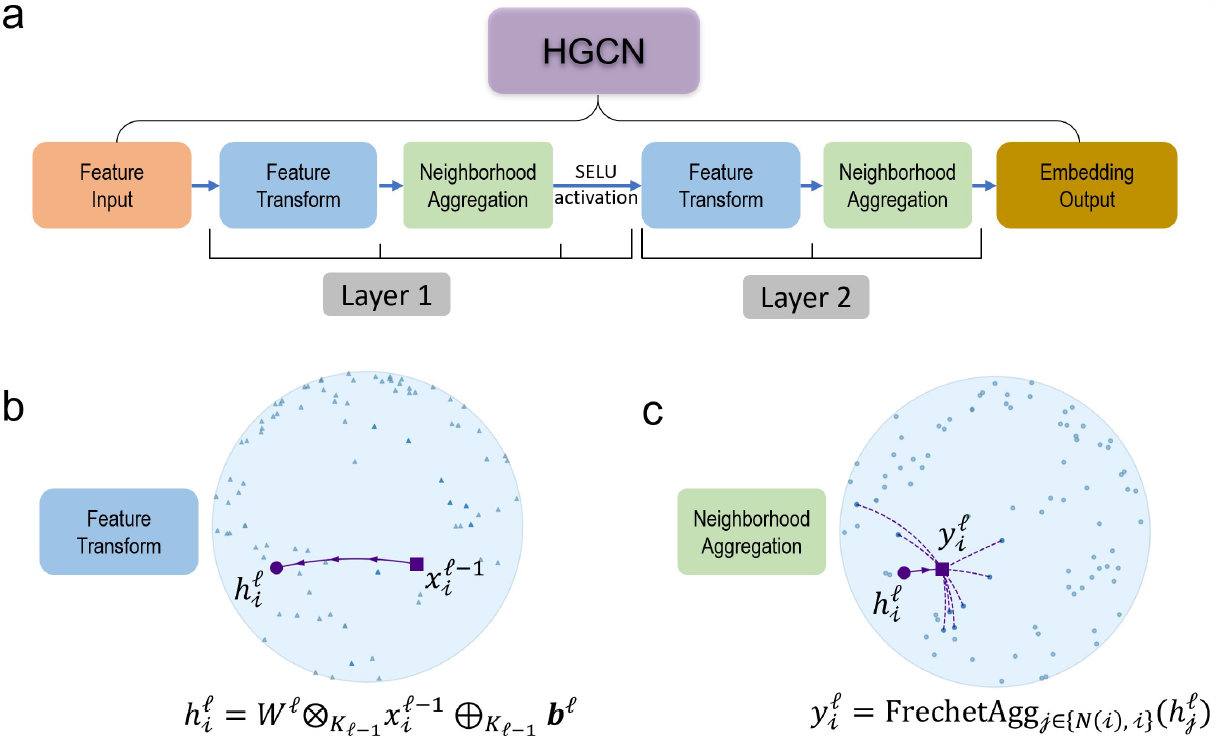
Architecture of the HGCN model. (a) Visualization of the two layers of the HGCN model. (b) Feature transform operation for an example node. (c) Fréchet aggregation for an example node. Plots use two dimensions for visualization purposes.

##### Input Features

In graph neural networks, node features represent additional information associated with each node in a graph. These attributes can be valuable for improving the performance of (H)GCNs in various tasks. In the absence of other relevant information, common choices for node features include one-hot identity vectors, node degrees, node position or coordinates, or the respective row of the adjacency matrix. Finding the proper balance of information density, simplicity, and bias is key to creating meaningful embeddings. Here, we used one-hot identity vectors.

##### Feature Transform

This operation combines hyperbolic matrix-vector multiplication and bias addition to implement a hyperbolic analogue of a fully connected linear layer:

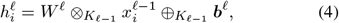

where 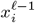 are the features of node *i* of the previous layer *ℓ* −1, and *W* ^*ℓ*^ and *b*^*ℓ*^ the weights and bias parameters of layer *ℓ*, respectively (Fig. 3b).

##### Fréchet Aggregation

An aggregation function captures local structure and features of the immediate neighborhood of each node. Such functions include weighted mean aggregation [45] and attention-based aggregation [46]. However, these methods are not trivial to incorporate in the hyperbolic space. The Fréchet mean, a generalized analogue of the Euclidean mean, does not have a closed form and, until recently, did not have a computationally efficient solver. The HGCN in [25] used the log/exp mapping to the tangent space to implement an attention mechanism to avoid the intractability of the Fréchet mean. However, recent work formulated a fully differentiable solution that outperforms the attention-based solutions [44]. Based on this, we substituted the attention-aggregation with the Fréchet mean aggregation (Fig. 3c):

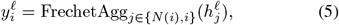

where *N*(*i*) denotes the neighbors of node *i*. In simple terms, the Fréchet mean is the point that minimizes the sum of distances (usually measured using a specific metric or distance function) from itself to all the data points in a set. It represents the “average” or “central” location within the dataset, considering the chosen distance metric.

##### Activation Function

The HGCN applies a hyperbolic non-linear activation function *σ*. We chose the ReLU function, implemented using the log/exp approach as in Eq. 3. The output of hidden layer was defined by:

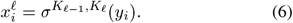

##### Weighted Loss Function

The HGCN models were trained using the Fermi-Dirac distribution as a decoder [27]. Under this distribution, the probability score of a link (or edge) between two nodes *i* and *j* at the output layer *L* is defined as:

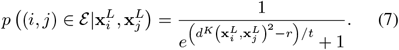

That is, an edge between two nodes is predicted to exist according to a sigmoidal transformation of the hyperbolic distance between the node embeddings. The hyperparameters *r* and *t* control the inflection point and steepness of the sigmoid function, and were set as *r* = 2 and *t* = 1.

The work in [25] used the Fermi-Dirac probabilities to train with cross-entropy loss for binary edge prediction. However here, in the case of MEG brain networks, we leveraged the continuous PLV metric to create a proxy value for the probability of connection by applying a min-max scaling of the original PLV values. With this formulation, the model was trained using mean square error of predicted and true probabilities, recovering information lost in the adjacency matrix thresholding.

## III Results

### A Parameter settings

Our implementation of HGCN contained two fully connected layers, comprising a hidden layer with output dimension 6, and a last layer with output dimension *D* (Fig. 3). It was trained using the Riemannian Adam optimizer [47] over 100 epochs with a single NVIDIA Tesla K80 GPU. Input features, batch size, and learning rate were set to one-hot identity, 1 and 0.02. We assessed the HGCN results using a model with fixed curvature *K* =*−*1 and a model with a learned curvature as a hyperparameter.

### B Link prediction results

To evaluate the effectiveness of hyperbolic graph embeddings in accurately representing MEG brain networks and determine the optimal hyperbolic embedding dimension, we performed link prediction experiments on the MEG brain networks [31]. Link prediction, a common downstream task within network science, aims to evaluate the capacity to identify missing connections within a network [13], [48].

We compared three well-established graph embedding models: our adapted HGCN [25], a shallow Poincaré embedding [27], and a graph convolutional network (GCN) with an identical architecture to the HGCN, albeit with flat curvature that reduces the Fréchet mean to a Euclidean mean [25]. While conventional GCNs generate MEG brain network representations in Euclidean space, both shallow Poincaré embedding and HGCN are able to encode high-dimensional brain networks into lower-dimensional hyperbolic manifolds, facilitating various downstream tasks, such as node or graph-level inference. However, the shallow Poincaré embedding approach has three critical limitations: 1) it cannot incorporate important node features when available, 2) it does not support inductive learning for graph inference tasks with unseen graph nodes, and 3) it is not applicable to large-scale graphs due to its inadequate scalability. Conversely, the HGCN model can effectively address the aforementioned issues and support an inductive graph embedding learning framework.

Mean averaged precision (MAP) results for the link prediction experiments with different embedding dimensions are shown in Table II. Note, the two HGCN models were trained with the Fermi-Dirac decoder [27] for link probability prediction, as shown in Eq. 7, using five-fold cross validation across participants. Specifically, the MEG brain networks for each participant, both before and after training, were consistently assigned to the same fold to eliminate any contamination of the validation sets.

**TABLE II.**
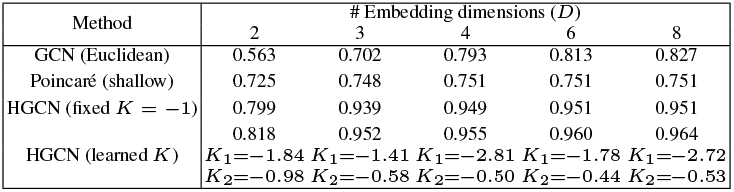
Comparison of mean averaged precision results for the link prediction task with different embedding dimensions (***D***) and three different graph embedding models.

The hyperbolic MEG brain network embeddings generated by the HGCN model achieved the highest link prediction performance, as measured by MAP, across all embedding dimensions (*D* = 2, 3, 4, 6, and 8), remarkably achieving a 0.952 MAP with merely three dimensions. In lower dimensions,the Euclidean GCN encountered constraints imposed by its geometry, resulting in inferior performance compared to the shallow Poincaré model. At dimensions *D* ≥ 4, the GCNs expressive learning power overcame its spatial limitations to outperform the shallow model, but still fell short against the HGCN. Notably, the HGCN with fixed curvature consistently trailed its learned curvature counterpart at all dimensions, as the latter possessed the adaptability to optimize curvature in each dimension for better fit. Interestingly, the HGCN with learned curvature exhibited a flattening trend after *D* = 3, suggesting that the need for strict hierarchy diminished as the space expanded.

### C SCD classification results

For SCD classification, we opted to apply the HGCN model with learned curvature and an embedding dimension *D* = 3. This decision stemmed from the preference for a simple low-dimensional model, given the diminishing gains at higher dimensions observed on Table II. Example hyperbolic embeddings for an HC and a SCD individual from the HGCN model, but with output embedding dimension *D* = 2 to ease visualization, are shown in Fig. 4. SCD classification relied on using the HGCN output embeddings as features in a traditional machine learning classifier to discriminate individuals with SCD from healthy controls [15].

**Fig. 4.**
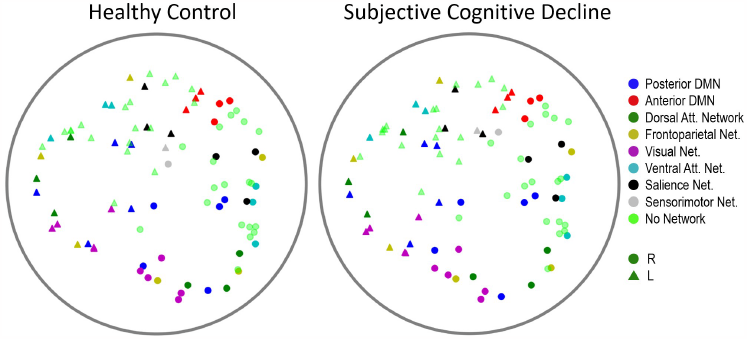
Examples of hyperbolic MEG brain network embeddings for a HC and a SCD participant. Individual points represent different brain regions, with the shape indicating hemisphere (circle for right hemisphere, triangle for left hemisphere), and color indicating subnetwork membership. To facilitate visualization, the output embedding dimension of the HGCN model had set to ***D = 2***.

To assess the quality of the HGCN hyperbolic embeddings, we considered three classification scenarios with features: (a) the entire raw PLV values contained in the adjacency matrices, (b) the distance from the hyperbolic center (i.e., radius) of the HGCN output embeddings of all ROIs, and (c) a combination of both sets of features. These features were fed to a support vector machine (SVM) classifier with a five-fold cross validation for supervised learning. Note, for this analysis, all features were limited to the pre-intervention MEG neural networks to remove any confounds from the cognitive training intervention.

The SCD classification results are shown in Fig. 5. The hyperbolic embedding features achieved higher performance compared to the PLV features, underscoring the distinct and robust information captured by the hyperbolic embeddings. In terms of the Macro F1 metric, the combination of hyperbolic embedding and PLV features improved performance. This suggests that these two feature types may contain partially complementary information, contributing positively to the classification task. However, this was not the case in the AUC-ROC metric, were the hyperbolic embeddings achieved a consistent score of 0.77, regardless of whether the PLV features were included or not. Overall, these results highlight the significance of the hyperbolic features as a potentially valuable diagnostic tool.

**Fig. 5.**
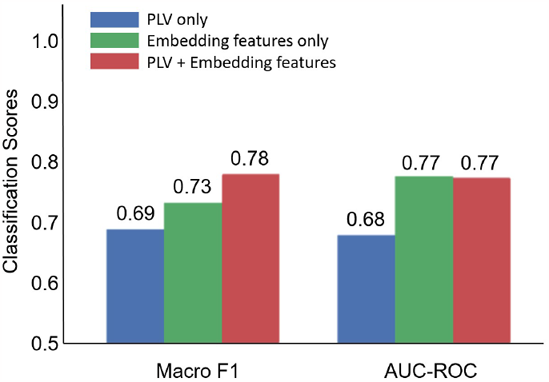
SCD classification performance based on PLV features, hyperboling embedding features, and their combination.

### D Brain subnetwork analysis

A disruption of brain connectivity in the SCD group could lead to subtle alterations that are encoded in the hyperbolic embeddings. To test this hypothesis, we studied the radius of the node embeddings, a metric that effectively proxies the hierarchical organization of the brain. This analysis was conducted by first averaging the radius of nodes within eight distinct functional subnetworks (pDMN, aDMN, DAN, FPN, VN, VAN, SN, SMN), and then investigating these values for SCD-related subnetwork effects.

Two-way repeated-measures ANCOVAs [49] with the within-subjects factor of time (pre-intervention/post-intervention) and between-subjects factor of diagnosis (SCD/HC), controlling for patient age, were conducted on the hyperbolic radius of each subnetwork, separately.

Statistical analysis used the SPSS Statistics software (Version 29) (Armonk, NY: IBM Corp.). Note, we did not use the training variable (trained/non-trained) as a factor in the ANCOVA. This is because exploratory analyses suggested that the groups pre-selected for training were not adequately randomly sampled in the distribution of age. This is relevant, since older age is a severity factor for SCD [50]. Levene’s test for equality of variances revealed there was a significant difference in variance of age between the trained and untrained groups (*F*(1, 88) = 4.532, *p* = .036). Therefore the assumption of homogeneity of variances was not met to qualify the use of the training variable in an ANCOVA. As a result, training groups were analyzed separately and age was included as a covariate so as to not promote erroneous effects.

The results of our statistical analysis indicate that the group receiving no cognitive training (n = 44) exhibited no significant effects after applying false discovery rate (FDR) corrections. In the group that underwent cognitive training (n = 46), three significant main effects of diagnosis survived FDR corrections (Fig. 6). Our findings are as follows: (i) a significant main effect of diagnosis on DAN hierarchy (*F* (1, 43) = 9.547, *p* = .0160, partial *η*^2^ = .182); (ii) on FPN hierarchy (*F* (1, 43) = 9.555, *p* = .0240, partial *η*^2^ = .182), and (iii) on VAN hierarchy (*F* (1, 43) = 8.563, *p* = .0133, partial *η*^2^ = .166). Inspecting estimated marginal means revealed that SCD individuals had a lower hyperbolic radius in the DAN (*M* = 1.786, *SE* = .036), FPN (*M* = 1.754, *SE* = .020), and VAN (*M* = 1.752, *SE* = .025) subnetworks compared to HC (*M* = 1.947, *SE* = .038; *M* = 1.844, *SE* = .021; and *M* = 1.859, *SE* = .026, respectively). It is worth noting that a lower cluster radius from the origin indicates a higher subnetwork hierarchy.

**Fig. 6.**
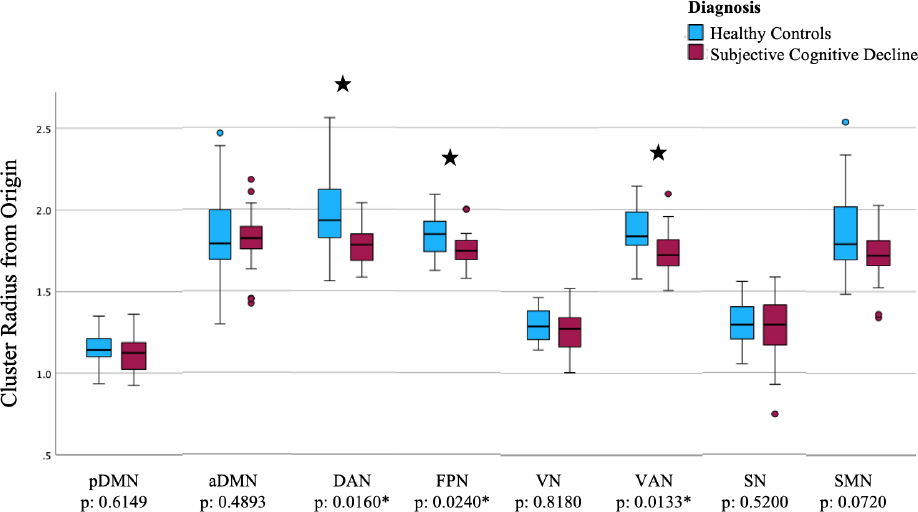
Hyperbolic radius of nodes belonging to each subnetwork for SCD and HC groups who received cognitive training derived from the final scan (pDMN: posterior default mode network. aDMN: anterior default mode network. DAN: dorsal attention network. FPN: frontoparietal network. VN: visual network. VAN: ventral attention network. SN: salience network. SMN: sensorimotor network).

### E Cognitive assessments

Our previous analysis, conducted for participants who received cognitive training, revealed that those experiencing SCD had a higher DAN, FPN, and VAN subnetwork hierarchy than healthy age-matched controls. Higher hierarchy of these significant subnetworks was correlated with poorer performance on multiple cognitive assessments. Specifically, we ran a Pearson’s bi-variate correlation [49] on the significant brain subnetwork embedding variables and cognitive assessment scores collected at post-intervention. The variables DAN radius, (*r* (81) = .223, *p* = .043) and VAN radius (*r* (81) =.245, *p* = .026) were significantly positively correlated with scores on the Rey-Osterrieth Complex Figure B copy test [51]. This indicates that higher hierarchy within the DAN and VAN subnetworks is significantly correlated with lower accuracy in reproducing a complex line drawing. Furthermore, the DAN radius was marginally negatively correlated with scores on the Geriatric Depression Scale (*r* (87) = -.206, *p* = .053). This suggests that higher DAN hierarchy was marginally associated with a greater number of depressive tendencies.

For a single subnetwork, the FPN, we found that lower hierarchy was correlated with an improvement on cognitive assessments over time. Specifically, we conducted Pearson’s bi-variate correlations between the radii of the DAN, FPN, and VAN subnetworks post-intervention and the change in cognitive assessment scores over time (post-intervention minus pre-intervention assessments). The radius of FPN was significantly negatively correlated with scores on the functional activities questionnaire (*r* (83) = -.261, *p* = .016). This suggests that lower FPN hierarchy was significantly correlated with improved performance on the functional activities questionnaire, meaning that participants reported having less impairment completing daily functional activities (i.e., preparing balanced meals, managing personal finances) over time. These results may suggest that the frontoparietal subnetwork is a key target when developing cognitive training interventions to elicit improvement over time.

## IV Discussion

We developed a hyperbolic MEG brain network embedding framework to study brain alterations in individuals with SCD. Our method automatically learns MEG brain network embeddings in a lower dimensional hyperbolic manifold, while maximally preserving the original topological properties of the brain networks. Furthermore, it effectively captures crucial hierarchical properties at both the node and subnetwork levels within the latent hyperbolic manifold, employing the hyperbolic radius as a proxy measure.

We compared our HGCN model, featuring both with fixed and learned curvature, against a Euclidean model and a shallow Poincaré model in the context of a link prediction task. Our HGCN model consistently outperformed all other models across all embedding dimensions, as evidenced in Table II. Notably, the HGCN model with learned curvature demonstrated higher accuracy than its fixed curvature counterpart. This finding suggests that the adaptability of curvature offers advantages in effectively modeling brain network data. Overall, the use of hyperbolic space within our model proves advantageous by preserving network distances and hierarchical information in the embeddings while minimizing distortion, making it particularly well-suited for handling complex “tree-like” neural data and large-scale brain graphs.

For SCD classification, we found that our HGCN embeddings markedly enhanced classifier performance compared to the functional connectivity (PLV) values in both Macro F1 and AUC-ROC. Using just five minutes of transformed resting-state neural data, we achieved Macro F1 of 0.73 and an AUC-ROC of 0.77, underscoring the promising diagnostic capabilities inherent in our approach. This outcome strongly implies that these embeddings contain intrinsic information related to brain network hierarchy affected by synaptic dysfunction, which can be effectively leveraged for machine-learning-based classification.

Our findings revealed brain network-level biomarkers for individuals experiencing SCD. In the cognitive training group, those with SCD displayed significantly elevated hierarchy within the DAN, FPN, and VAN subnetworks when compared to their healthy counterparts. Notably, this increased hierarchy in these subnetworks was associated with poorer cognitive performance across all participants. This discovery holds particular significance given that SCD represents a potential precursor to AD, a condition that cannot be early diagnosed through standardized cognitive assessments.

Although largely associated with AD, subjective cognitive decline is a heterogeneous condition. As many as 60 percent of individuals with SCD progress to MCI or AD over a 15 year period [52]. This also means that 40 percent of older adults with SCD remain stable or may experience related conditions. It is important to recognize that SCD is linked not only to AD but also to various non-AD forms of dementia, Parkinson’s disease, cerebrovascular disease, psychiatric disorders, medication or substance effects, and even normal aging [3]. To ensure the validity of our study, we meticulously excluded potential confounding factors such as psychiatric and neurological diseases, medication, and substance use. Our cohort was carefully selected based on multiple criteria that increase the likelihood of AD development in individuals with SCD. These criteria include self-reported cognitive concerns primarily related to memory, an age above 60 years at the onset of SCD, an onset occurring within the last 5 years, and cognitive decline confirmed by a reliable informant [6].

Our study revealed ROI connectivity dysfunction in the DAN, VAN, and FPN subnetworks in participants with SCD compared to healthy controls. These findings align with the current body of literature on AD as a progressive neurodegenerative condition. Previous research has already established that both the VAN and DAN networks [53]–[55], and the FPN [56], [57], exhibit altered functional connectivity in AD. The attention networks are specialized for distinct processes such as the detection of unexpected but behaviorally relevant stimuli (VAN) and top-down controlled attention (DAN). The VAN and DAN can flexibly interact depending on task demands, with frontal regions like the inferior and middle frontal gyrus playing a moderating role [58]. The frontoparietal network regulates the moment-to-moment decisions involved in the planning and execution of goal directed behavior (i.e. processes such as working memory, inhibitory control, capacity to focus and screen out interfering information, the ability to formulate an action plan) [59].

At prodromal stages, alterations in the DAN have been consistently observed in patients with MCI [19], [60], [61] and SCD [19], [62], [63]. These alterations have been linked to behavioral deficits in top-down attentional control [61]. Furthermore, a MEG resting-state study examined functional connectivity alterations between healthy controls, individuals with SCD, and MCI patients [19]. This study reported both SCD individuals and MCI patients displayed a similar spatial pattern of functional connectivity (FC) alterations with evidence of progressive deterioration. This pattern was characterized by a hyper-synchronized anterior network and a hypo-synchronized posterior network.

Although our findings exist at the brain subnetwork level and lack ROI-to-ROI level specificity, a review of relevant literature offers further insights. When comparing clinical participants (SCD and MCI) to healthy controls, noteworthy alterations emerge in the bilateral *supramarginal gyrus* and *right angular gyrus*. These regions exhibit reduced functional connectivity with medial temporal areas (including the hippocampus and parahippocampal gyrus) as well as several occipital areas [19]. These ROIs, integral to the frontoparietal and ventral attention networks, may play a pivotal role in network-level disruptions at preclinical stages of AD. Moreover, the *insula*, a pivotal component of the ventral attention network, exhibits connectivity alterations during this preclinical stage [63]. In one study, SCD participants displayed lower insula activation during a task involving switching between imagination and temporal decision-making, suggesting impaired control over proper engagement of the DMN and FPN, which govern executive control functions [64]. In another study, SCD participants demonstrated weakened connectivity between the DMN and FPN [65]. These connectivity findings are conceptually consistent with the pattern of reduced gray matter density seen in both SCD and MCI within the bilateral medial temporal, frontotemporal, and neocortical areas. This gray matter density reduction in medial temporal regions inversely correlated with the cognitive complaint index [66], establishing a neural basis for these cognitive complaints.

Our study did not find any hierarchy changes in the anterior or posterior DMN. Even though alterations in DMN connectivity have been extensively documented in individuals with MCI and AD [67]–[69], research on SCD individuals has produced mixed results [19], [63], [65], [70], [71]. Some studies have reported hypo-connectivity in the pDMN [19] and cDMN [65], while some study has found hyper-connectivity in the DMN as a whole [63]. These discrepancies may stem from variations in the criteria used for defining regions of interest (ROIs) or the segmentation of the Default Mode Network into its constituent parts. Additionally, the mixed results may be attributed to concurrent hyper- and hypo-connectivity within the DMN, which are observed in AD and MCI groups [67] and may manifest at the preclinical stage.

Last, our ANCOVA analysis revealed no significant effects directly attributable to the cognitive intervention. This absence of findings may be attributed to the possibility that any changes induced by the cognitive training were exceedingly subtle to be captured by our modeling. In terms of limitations, our study was confined to a single modality, relying solely on MEG data. While MEG is well-suited for assessing synaptic dysfunction, it lacks precise anatomical specificity. Additionally, our study produced significant findings with a relatively modest sample size. Subsequent investigations should prioritize the inclusion of a larger participant pool to augment the statistical reliability of our results. Moreover, incorporating multi-modal MEG and MRI data could extend and refine our methods, providing a more comprehensive understanding of brain network alterations in the context of cognitive decline. Finally, the inclusion of amyloid biomarkers could provide invaluable insights into the underlying mechanisms driving the alterations in brain networks, but unfortunately such information was not available in our cohort.

## V Conclusion

We developed a novel hyperbolic brain network embedding method to transform complex MEG brain networks into low-dimensional hyperbolic embeddings. Our method method effectively preserves network geometric distances and hierarchical information with low distortion, appropriately handles large-scale graphs, and utilizes inductive learning for graph interference tasks with unseen nodes. With this approach, we examined alterations in brain networks among individuals with SCD in comparison to healthy controls. Our HGCN model with learned curvature outperformed all other tested models on a link prediction task. Furthermore, when compared to PLV connectivity values, our HGCN embeddings significantly enhanced classifier performance in distinguishing between SCD and healthy controls, highlighting the diagnostic potential of transformed resting-state neural data.

Notably, we introduced a unique metric — the radius of the node embeddings — which effectively proxies the hierarchical organization of the brain. We leveraged this metric to characterize subtle hierarchical organization changes of various brain subnetworks in the SCD population. We found increased hierarchy, indicated by reduced hyperbolic radii, in the DAN, VAN, and FPN subnetworks for participants experiencing SCD. These findings align with existing evidence of dysfunction in these subnetworks observed in SCD [19], [62], [63], and MCI and AD patients [55], [60], [61]. Furthermore, these three subnetworks were correlated with diminished performance on multiple cognitive assessments.

In conclusion, our study contributes a valuable framework for exploring brain network alterations associated with cognitive decline. The utilization of hyperbolic embeddings and the novel hierarchical metric provide fresh insights into the intricate dynamics of brain networks in the context of SCD, offering potential avenues for future research and clinical applications. We share code and data to reproduce this work here: https://github.com/ColeSBaker/hyperBrain.

## References

[1] C. Qiu, M. Kivipelto, and E. Von Strauss, “Epidemiology of alzheimer’s disease: occurrence, determinants, and strategies toward intervention,” Dialogues in clinical neuroscience, 2022.

[2] K. B. Rajan, J. Weuve, L. L. Barnes, E. A. McAninch, R. S. Wilson, and D. A. Evans, “Population estimate of people with clinical alzheimer’s disease and mild cognitive impairment in the united states (2020–2060),” Alzheimer’s & dementia, vol. 17, no. 12, pp. 1966–1975, 2021.

[3] F. Jessen, R. E. Amariglio, M. Van Boxtel, M. Breteler, M. Ceccaldi, G. Chételat, B. Dubois, C. Dufouil, K. A. Ellis, W. M. Van Der Flier et al., “A conceptual framework for research on subjective cognitive decline in preclinical alzheimer’s disease,” Alzheimer’s & dementia, vol. 10, no. 6, pp. 844–852, 2014.

[4] C. Morrison, M. Dadar, N. Shafiee, S. Villeneuve, D. L. Collins, A. D. N. Initiative et al., “Regional brain atrophy and cognitive decline depend on definition of subjective cognitive decline,” NeuroImage: Clinical, vol. 33, p. 102923, 2022.

[5] H. Lin, J. Jiang, Z. Li, C. Sheng, W. Du, X. Li, and Y. Han, “Identification of subjective cognitive decline due to alzheimer’s disease using multimodal mri combining with machine learning,” Cerebral Cortex, 2022.

[6] L. A. Rabin, C. M. Smart, and R. E. Amariglio, “Subjective cognitive decline in preclinical alzheimer’s disease,” Annual review of clinical psychology, vol. 13, pp. 369–396, 2017.

[7] B. Reisberg, “The pre-mild cognitive impairment, subjective cognitive impairment stage of alzheimer’s disease,” Alzheimer’s I & Dementia, 2008.

[8] J. Hardy and D. J. Selkoe, “The amyloid hypothesis of alzheimer’s disease: progress and problems on the road to therapeutics,” science, vol. 297, no. 5580, pp. 353–356, 2002.

[9] V. Garcia-Marin, L. Blazquez-Llorca, J.-R. Rodriguez, S. Boluda, G. Muntane, I. Ferrer, and J. DeFelipe, “Diminished perisomatic gabaergic terminals on cortical neurons adjacent to amyloid plaques,” Frontiers in neuroanatomy, vol. 3, p. 1102, 2009.

[10] T. Taniguchi, T. Kawamata, H. Mukai, H. Hasegawa, T. Isagawa, M. Yasuda, T. Hashimoto, A. Terashima, M. Nakai, Y. Ono et al., “Phosphorylation of tau is regulated by pkn,” Journal of Biological Chemistry, vol. 276, no. 13, pp. 10025–10031, 2001.

[11] F. Maestú, J.-M. Peña, P. Garcés, S. González, R. Bajo, A. Bagic, P. Cuesta, M. Funke, J. P. Mäkelä, E. Menasalvas et al., “A multicenter study of the early detection of synaptic dysfunction in mild cognitive impairment using magnetoencephalography-derived functional connectivity,” NeuroImage: Clinical, vol. 9, pp. 103–109, 2015.

[12] K. Wang, M. Liang, L. Wang, L. Tian, X. Zhang, K. Li, and T. Jiang, “Altered functional connectivity in early alzheimer’s disease: A restingstate fmri study,” Human brain mapping, vol. 28, no. 10, pp. 967–978, 2007.

[13] M. Xu, Z. Wang, H. Zhang, D. Pantazis, H. Wang, and Q. Li, “A new graph gaussian embedding method for analyzing the effects of cognitive training,” PLoS computational biology, vol. 16, no. 9, p. e1008186, 2020.

[14] Y. Ding, J. H. Sohn, M. G. Kawczynski, H. Trivedi, R. Harnish, N. W. Jenkins, D. Lituiev, T. P. Copeland, M. S. Aboian, C. Mari Aparici et al., “A deep learning model to predict a diagnosis of alzheimer disease by using 18f-fdg pet of the brain,” Radiology, vol. 290, no. 2, pp. 456–464, 2019.

[15] M. Xu, D. L. Sanz, P. Garces, F. Maestu, Q. Li, and D. Pantazis, “A graph gaussian embedding method for predicting alzheimer’s disease progression with meg brain networks,” IEEE Transactions on Biomedical Engineering, vol. 68, no. 5, pp. 1579–1588, 2021.

[16] F. Maestú, P. Cuesta, O. Hasan, A. Fernandéz, M. Funke, and P. E. Schulz, “The importance of the validation of m/eeg with current biomarkers in alzheimer’s disease,” Frontiers in human neuroscience, vol. 13, p. 17, 2019.

[17] S. Baillet, “Magnetoencephalography for brain electrophysiology and imaging,” Nature neuroscience, vol. 20, no. 3, pp. 327–339, 2017.

[18] D. Pantazis, M. Fang, S. Qin, Y. Mohsenzadeh, Q. Li, and R. M. Cichy, “Decoding the orientation of contrast edges from meg evoked and induced responses,” NeuroImage, vol. 180, pp. 267–279, 2018.

[19] D. López-Sanz, R. Bruña, P. Garcés, M. C. Martín-Buro, S. Walter, M. L. Delgado, M. Montenegro, R. López Higes, A. Marcos, and F. Maestú, “Functional connectivity disruption in subjective cognitive decline and mild cognitive impairment: a common pattern of alterations,” Frontiers in aging neuroscience, vol. 9, p. 109, 2017.

[20] C.-H. Cheng, P.-N. Wang, H.-F. Mao, and F.-J. Hsiao, “Subjective cognitive decline detected by the oscillatory connectivity in the default mode network: a magnetoencephalographic study,” Aging (Albany NY), vol. 12, no. 4, p. 3911, 2020.

[21] G. Rosenthal, F. Vása, A. Griffa, P. Hagmann, E. Amico, J. Goñi, G. Avidan, and O. Sporns, “Mapping higher-order relations between brain structure and function with embedded vector representations of connectomes,” Nature communications, vol. 9, no. 1, pp. 1–12, 2018.

[22] A. Bojchevski and S. Gunnemann, “Deep gaussian embedding of graphs: Unsupervised inductive learning via ranking,” arXiv preprint arXiv:1707.03815, 2017.

[23] M. Xu, “Understanding graph embedding methods and their applications,” SIAM Review, vol. 63, no. 4, pp. 825–853, 2021.

[24] A. Gu, F. Sala, B. Gunel, and C. Ré, “Learning mixed-curvature representations in product spaces,” in International Conference on Learning Representations, 2018.

[25] I. Chami, Z. Ying, C. Ré, and J. Leskovec, “Hyperbolic graph convolutional neural networks,” Advances in neural information processing systems, vol. 32, 2019.

[26] F. Di Giovanni, G. Luise, and M. Bronstein, “Heterogeneous manifolds for curvature-aware graph embedding,” arXiv preprint arXiv:2202.01185, 2022.

[27] M. Nickel and D. Kiela, “Poincaré embeddings for learning hierarchical representations,” Advances in neural information processing systems, vol. 30, 2017.

[28] M. Nickel and D. Kiela, “Learning continuous hierarchies in the lorentz model of hyperbolic geometry,” in International Conference on Machine Learning. PMLR, 2018, pp. 3779–3788.

[29] Q. Liu, M. Nickel, and D. Kiela, “Hyperbolic graph neural networks,” Advances in Neural Information Processing Systems, vol. 32, 2019.

[30] W. Peng, T. Varanka, A. Mostafa, H. Shi, and G. Zhao, “Hyperbolic deep neural networks: A survey,” IEEE Transactions on pattern analysis and machine intelligence, vol. 44, no. 12, pp. 10 023–10 044, 2021.

[31] Suárez-Méndez, R. Bruña, D. López-Sanz, P. Montejo, M. Montenegro-Peña, M. L. Delgado-Losada, A. Marcos Dolado, R. López-Higes, and F. Maestú, “Cognitive training modulates brain hypersynchrony in a population at risk for alzheimer’s disease,” Journal of Alzheimer’s Disease, vol. 86, no. 3, pp. 1185–1199, 2022.

[32] S. Taulu and R. Hari, “Removal of magnetoencephalographic artifacts with temporal signal-space separation: Demonstration with single-trial auditory-evoked responses,” Human Brain Mapping, 2009.

[33] N. T.-M. N, “Automated anatomical labeling of activations in spm using a macroscopic anatomical parcellation of the mni mri single-subject brain.” Neuroimage, 2001.

[34] G. Nolte, “The magnetic lead field theorem in the quasi-static approximation and its use for magnetoenchephalography forward calculation in realistic volume conductors.” Phys Med Biol, 2003.

[35] V. Veen, “Localization of brain electrical activity via linearly constrained minimum variance spectral filtering.” IEEE Trans Biomed Eng, 1997.

[36] Z. Marton-Alper, A. Markus, M. Nevat, R. Bennet, and S. G. Shamay-Tsoory, “Differential contribution of between and within-brain coupling to movement synchronization,” Human Brain Mapping, 2023.

[37] M. Rubinov and E. Bullmore, “Schizophrenia and abnormal brain network hubs,” Dialogues in clinical neuroscience, 2022.

[38] W. Shirer, “Decoding subject-driven cognitive states with whole-brain connectivity patterns.” Cerebral Cortex, 2011.

[39] P. Boveroux, “Breakdown of within- and between-network resting state functional magnetic resonance imaging connectivity during propofolinduced loss of consciousness.” Anesthesiology, 2010.

[40] B. He, “Breakdown of functional connectivity in frontoparietal networks underlies behavioral deficits in spatial neglect.” Neuron, 2007.

[41] A. Janes, “Insula–dorsal anterior cingulate cortex coupling is associated with enhanced brain reactivity to smoking cues.” Neuropsychopharmacology, 2010.

[42] O.-E. Ganea and G. Bécigneul, “Hyperbolic neural networks,” NIPS ‘18, 2018.

[43] R. Shimizu, “Hyperbolic neural networks ++,” ICLR 2021, 2021.

[44] A. Lou, “Differentiating through the frechet mean,” Proceedings of the 37th International Conference on Machine Learning, 2020.

[45] F. Zhang, Z. Liu, F. Xiong, J. Su, and H. Qiao, “Wagnn: A weighted aggregation graph neural network for robot skill learning,” Robotics and Autonomous Systems, vol. 130, p. 103555, 2020.

[46] P. Velickovi’c, G. Cucurull, A. Casanova, A. Romero, P. Li’o, and Y. Bengio, “Graph attention networks,” ICLR, 2018.

[47] M. Kochurov, R. Karimov, and S. Kozlukov, “Geoopt: Riemannian optimization in pytorch,” 2020.

[48] M. Zhang and Y. Chen, “Link prediction based on graph neural networks,” Advances in neural information processing systems, vol. 31, 2018.

[49] A. Field, Discovering statistics using IBM SPSS statistics. sage, 2013.

[50] C. Wen, H. Hu, Y.-N. Ou, Y.-L. Bi, Y.-H. Ma, L. Tan, and J.-T. Yu, “Risk factors for subjective cognitive decline: the cable study,” Translational psychiatry, vol. 11, no. 1, pp. 1–9, 2021.

[51] S. Luzzi, M. Pesallaccia, K. Fabi, M. Muti, G. Viticchi, L. Provinciali, and M. Piccirilli, “Non-verbal memory measured by rey–osterrieth complex figure b: normative data,” Neurological Sciences, vol. 32, pp. 1081–1089, 2011.

[52] B. Reisberg and S. Gauthier, “Current evidence for subjective cognitive impairment (sci) as the pre-mild cognitive impairment (mci) stage of subsequently manifest alzheimer’s disease,” International psychogeriatrics, vol. 20, no. 1, pp. 1–16, 2008.

[53] X. Wu, R. Li, A. S. Fleisher, E. M. Reiman, X. Guan, Y. Zhang, K. Chen, and L. Yao, “Altered default mode network connectivity in alzheimer’s disease—a resting functional mri and bayesian network study,” Human brain mapping, vol. 32, no. 11, pp. 1868–1881, 2011.

[54] R. Li, X. Wu, A. S. Fleisher, E. M. Reiman, K. Chen, and L. Yao, “Attention-related networks in alzheimer’s disease: A resting functional mri study,” Human brain mapping, vol. 33, no. 5, pp. 1076–1088, 2012.

[55] Wang, J. Liu, Z. Wang, P. Sun, K. Li, and P. Liang, “Dysfunctional interactions between the default mode network and the dorsal attention network in subtypes of amnestic mild cognitive impairment,” Aging (Albany NY), vol. 11, no. 20, p. 9147, 2019.

[56] Q. Zhao, H. Lu, H. Metmer, W. X. Li, and J. Lu, “Evaluating functional connectivity of executive control network and frontoparietal network in alzheimer’s disease,” Brain research, vol. 1678, pp. 262–272, 2018.

[57] Z. Wang, K. Qiao, G. Chen, D. Sui, H.-M. Dong, Y.-S. Wang, H.-J. Li, Lu, X.-N. Zuo, and Y. Han, “Functional connectivity changes across the spectrum of subjective cognitive decline, amnestic mild cognitive impairment and alzheimer’s disease,” Frontiers in neuroinformatics, vol. 13, p. 26, 2019.

[58] S. Vossel, J. J. Geng, and G. R. Fink, “Dorsal and ventral attention systems: distinct neural circuits but collaborative roles,” The Neuroscientist, vol. 20, no. 2, pp. 150–159, 2014.

[59] E. Mezzacappa, “Executive function,” in Encyclopedia of Adolescence, B. B. Brown and M. J. Prinstein, Eds. San Diego: Academic Press, 2011, pp. 142–150. [Online]. Available: https://www.sciencedirect.com/science/article/pii/B9780123739513000168

[60] S. Qian, Z. Zhang, B. Li, and G. Sun, “Functional-structural degeneration in dorsal and ventral attention systems for alzheimer’s disease, amnestic mild cognitive impairment,” Brain imaging and behavior, vol. 9, pp. 790–800, 2015.

[61] Z. Zhang, H. Zheng, K. Liang, H. Wang, S. Kong, J. Hu, F. Wu, and G. Sun, “Functional degeneration in dorsal and ventral attention systems in amnestic mild cognitive impairment and alzheimer’s disease: an fmri study,” Neuroscience letters, vol. 585, pp. 160–165, 2015.

[62] H. Wu, Y. Song, X. Yang, S. Chen, H. Ge, Z. Yan, W. Qi, Q. Yuan, X. Liang, X. Lin et al., “Functional and structural alterations of dorsal attention network in preclinical and early-stage alzheimer’s disease,” CNS Neuroscience & Therapeutics, vol. 29, no. 6, pp. 1512–1524, 2023.

[63] D. Lee, J. Y. Park, and W. J. Kim, “Altered functional connectivity of the default mode and dorsal attention network in subjective cognitive decline,” Journal of Psychiatric Research, vol. 159, pp. 165–171, 2023.

[64] X. Hu, F. Uhle, K. Fliessbach, M. Wagner, Y. Han, B. Weber, and F. Jessen, “Reduced future-oriented decision making in individuals with subjective cognitive decline: A functional mri study,” Alzheimer’s & Dementia: Diagnosis, Assessment & Disease Monitoring, vol. 6, pp. 222–231, 2017.

[65] Y.-C. Wei, Y.-C. Kung, W.-Y. Huang, C. Lin, Y.-L. Chen, C.-K. Chen, Y.-C. Shyu, and C.-P. Lin, “Functional connectivity dynamics altered of the resting brain in subjective cognitive decline,” Frontiers in Aging Neuroscience, vol. 14, p. 817137, 2022.

[66] A. Saykin, H. Wishart, L. Rabin, R. Santulli, L. Flashman, J. West,T. McHugh, and A. Mamourian, “Older adults with cognitive complaints show brain atrophy similar to that of amnestic mci,” Neurology, vol. 67, no. 5, pp. 834–842, 2006.

[67] A. Badhwar, A. Tam, C. Dansereau, P. Orban, F. Hoffstaedter, and P. Bellec, “Resting-state network dysfunction in alzheimer’s disease: a systematic review and meta-analysis,” Alzheimer’s & Dementia: Diagnosis, Assessment & Disease Monitoring, vol. 8, pp. 73–85, 2017.

[68] C. Sorg, V. Riedl, M. Muhlau, V. D. Calhoun, T. Eichele, L. Läer, A. Drzezga, H. Förstl, A. Kurz, C. Zimmer et al., “Selective changes of resting-state networks in individuals at risk for alzheimer’s disease,” Proceedings of the National Academy of Sciences, vol. 104, no. 47, pp. 18 760–18 765, 2007.

[69] F.-J. Hsiao, Y.-J. Wang, S.-H. Yan, W.-T. Chen, and Y.-Y. Lin, “Altered oscillation and synchronization of default-mode network activity in mild alzheimer’s disease compared to mild cognitive impairment: an electrophysiological study,” PloS one, vol. 8, no. 7, p. e68792, 2013.

[70] C. Xue, W. Qi, Q. Yuan, G. Hu, H. Ge, J. Rao, C. Xiao, and J. Chen, “Disrupted dynamic functional connectivity in distinguishing subjective cognitive decline and amnestic mild cognitive impairment based on the triple-network model,” Frontiers in Aging Neuroscience, vol. 13, p. 711009, 2021.

[71] R. P. Viviano and J. S. Damoiseaux, “Longitudinal change in hippocampal and dorsal anterior insulae functional connectivity in subjective cognitive decline,” Alzheimer’s Research & Therapy, vol. 13, no. 1, p. 108, 2021.

