## Supplementary Material for "Hyperbolic graph embedding of MEG brain networks to study brain alterations in individuals with subjective cognitive decline" (from another version of this preprint: version 2, 2023-10-28)

### I. Assignment of brain regions to subnetworks

Atlases delineating ROIs within brain subnetworks deviate significantly from structural atlases like the AAL because they are based on patterns of communication instead of physical structures in the brain. To assign brain regions to distinct subnetworks, we conducted a comprehensive evaluation of various subnetwork studies [1]–[4]. We considered eight distinct brain subnetworks: posterior default mode network (pDMN), anterior default mode network (aDMN), dorsal attention network (DAN), frontoparietal network (FPN), visual network (VN), ventral attention network (VAN), salience network (SN), and sensorimotor network (SMN). We then devised a procedure that considered the spatial proximity between each AAL anatomical region and the central coordinates of different functional ROIs.

Each brain subnetwork contained anywhere between 4 to 12 functional ROIs. To represent each functional ROI, we used its 3D center coordinates. In order to assign AAL anatomical ROIs to these subnetwork functional ROIs, we employed a straightforward technique that considered the spatial proximity between these regions. Specifically, for each source $A_k$ located within an anatomical ROI, we computed the distance $\mathrm{dist}(C_{A_k}, C_{F_j})$ to the center of the corresponding functional ROI $C_{F_j}$. We then defined an inclusion score, denoting the percentage of sources for which the distance fell within a defined threshold $r$, as follows:

$$P_{A,F} = \frac{1}{N_A} \sum_{N_A} \sum_{N_F} \begin{cases} 1, & \text{if } \mathrm{dist}(C_{A_k}, C_{F_j}) < r \\ 0, & \text{otherwise} \end{cases} \quad (1)$$

where $A$, $N_A$, and $A_k$ are an ROI in $AAL$, the number of sources in $A$ and the $kth$ source in $A$. $F$ is a subnetwork, $F_j$ is the $jth$ functional ROI belonging to $F$, $N_F$ is the number of functional ROIs in $F$, $P_{A,F}$ is the percentage score of $A$ belonging to $F$. Note that an average is only taken over the number $N_A$, not $N_F$, because the score calculates the percentage of *all* nodes in $A$ that are near *any* ROI in $F$.

The distance threshold $r$ was set to 1.5 cm. Last, the inclusion threshold $t$ was selected such that $A$ is assigned to $F$ if $P_{A,F} > t$. Anatomical ROIs were allowed to be assigned to multiple subnetworks. The value of $t$ was set at 0.3, as it limited all subnetworks to a reasonable while ensuring that regions known to belong to specific subnetworks were correctly assigned.

The list of anatomical ROIs belonging to each subnetwork are detailed in the tables below.

### References


[1] W. Shirer, "Decoding subject-driven cognitive states with whole-brain connectivity patterns." *Cerebral Cortex*, 2011.

[2] P. Boveroux, "Breakdown of within- and between-network resting state functional magnetic resonance imaging connectivity during propofol-induced loss of consciousness." *Anesthesiology*, 2010.

[3] B. He, "Breakdown of functional connectivity in frontoparietal networks underlies behavioral deficits in spatial neglect." *Neuron*, 2007.

[4] A. Janes, "Insula–dorsal anterior cingulate cortex coupling is associated with enhanced brain reactivity to smoking cues." *Neuropsychopharmacology*, 2010.

| Posterior Default Mode Network (pDMN) | |
|---|---|
| Names | Nicknames |
| ['Left Cingulate gyrus, Posterior part'] | ['lPCC'] |
| ['Right Cingulate gyrus, Posterior part'] | ['rPCC'] |
| ['Left Hippocampus'] | ['lHip'] |
| ['Right Hippocampus'] | ['rHip'] |
| ['Left Parahippocampus'] | ['lParahip'] |
| ['Right Parahippocampus'] | ['rParahip'] |
| ['Left Angular gyrus'] | ['lAng'] |
| ['Right Angular gyrus'] | ['rAng'] |
| ['Left Precuneus'] | ['lPrecu'] |
| ['Right Precuneus'] | ['rPrecu'] |

Fig. S1: pDMN ROI Assignment

| Anterior Default Mode Network (aDMN) | |
|---|---|
| Names | Nicknames |
| ['Left Superior Frontal gyrus, Medial'] | ['lSFGm'] |
| ['Right Superior Frontal gyrus, Medial'] | ['rSFGm'] |
| ['Left Superior Frontal gyrus, Medial Orbital'] | ['lSFGmo'] |
| ['Right Superior Frontal gyrus, Medial Orbital'] | ['rSFGmo'] |
| ['Left Cingulate gyrus, Anterior part'] | ['lACC'] |
| ['Right Cingulate gyrus, Anterior part'] | ['rACC'] |

Fig. S2: aDMN ROI Assignment

| Dorsal Attention Network (DAN) | |
|---|---|
| Names | Nicknames |
| ['Left Precentral gyrus'] | ['lPreCG'] |
| ['Right Precentral gyrus'] | ['rPreCG'] |
| ['Left Inferior Parietal gyrus'] | ['lIPG'] |
| ['Right Inferior Parietal gyrus'] | ['rIPG'] |
| ['Left Superior Parietal gyrus'] | ['lSPG'] |
| ['Right Superior Parietal gyrus'] | ['rSPG'] |

Fig. S3: DAN ROI Assignment

| Fronto-Pariental Network (FPN) | |
|---|---|
| Names | Nicknames |
| ['Left Inferior Frontal gyrus, Triangular'] | ['lIFGt'] |
| ['Right Inferior Frontal gyrus, Triangular'] | ['rIFGt'] |
| ['Left Superior Occipital lobe'] | ['lSOccL'] |
| ['Right Superior Occipital lobe'] | ['rSOccL'] |
| ['Left Inferior Parietal gyrus'] | ['lIPG'] |
| ['Right Inferior Parietal gyrus'] | ['rIPG'] |
| ['Left Supramarginal gyrus'] | ['lSMG'] |
| ['Right Supramarginal gyrus'] | ['rSMG'] |
| ['Left Angular gyrus'] | ['lAng'] |
| ['Right Angular gyrus'] | ['rAng'] |

Fig. S4: FPN ROI Assignment

| Visual Network (VN) | |
|---|---|
| Names | Nicknames |
| ['Left Calcarine fissure'] | ['lCalc'] |
| ['Right Calcarine fissure'] | ['rCalc'] |
| ['Left Cuneus'] | ['lCu'] |
| ['Right Cuneus'] | ['rCu'] |
| ['Left Lingual gyrus'] | ['lLingual'] |
| ['Right Lingual gyrus'] | ['rLingual'] |
| ['Left Middle Occipital lobe'] | ['lMOccL'] |
| ['Right Middle Occipital lobe'] | ['rMOccL'] |
| ['Left Inferior Occipital lobe'] | ['lIOccL'] |
| ['Right Inferior Occipital lobe'] | ['rIOccl'] |

Fig. S5: VN ROI Assignment

| Ventral Attention Network (VAN) | |
|---|---|
| Names | Nicknames |
| ['Left Precentral gyrus'] | ['lPreCG'] |
| ['Right Precentral gyrus'] | ['rPreCG'] |
| ['Left Inferior Frontal gyrus, Opercular'] | ['lIFGo'] |
| ['Right Inferior Frontal gyrus, Opercular'] | ['rIFGor'] |
| ['Left Inferior Frontal gyrus, Triangular'] | ['lIFGt'] |
| ['Right Inferior Frontal gyrus, Triangular'] | ['rIFGt'] |
| ['Left Insula'] | ['lInsula'] |
| ['Right Insula'] | ['rInsula'] |
| ['Left Supramarginal gyrus'] | ['lSMG'] |
| ['Right Supramarginal gyrus'] | ['rSMG'] |
| ['Left Angular gyrus'] | ['lAng'] |
| ['Right Angular gyrus'] | ['rAng'] |
| ['Left Middle temporal gyrus'] | ['lMTG'] |
| ['Right Middle temporal gyrus'] | ['rMTG'] |

Fig. S6: VAN ROI Assignment

| Salience Network (SN) | |
|---|---|
| Names | Nicknames |
| ['Left Inferior Frontal gyrus, Orbital'] | ['lIFGo'] |
| ['Right Inferior Frontal gyrus, Orbital'] | ['rIFGo'] |
| ['Left Olfactory cortex'] | ['lOlfC'] |
| ['Right Olfactory cortex'] | ['rOlfC'] |
| ['Left Cingulate gyrus, Anterior part'] | ['lACC'] |
| ['Right Cingulate gyrus, Anterior part'] | ['rACC'] |
| ['Left Hippocampus'] | ['lHip'] |
| ['Right Hippocampus'] | ['rHip'] |
| ['Left Parahippocampus'] | ['lParahip'] |
| ['Right Parahippocampus'] | ['rParahip'] |
| ['Left Amygdala'] | ['lAmyg'] |
| ['Right Amygdala'] | ['rAmyg'] |

Fig. S7: SN ROI Assignment

| Sensori-Motor Network (SMN) | |
|---|---|
| Names | Nicknames |
| ['Left Supplementary Motor area'] | ['lMotor'] |
| ['Right Supplementary Motor area'] | ['rMotor'] |

Fig. S8: SMN ROI Assignment
